# Sexual dimorphism of MASLD-driven bone loss

**DOI:** 10.1101/2024.11.25.625246

**Authors:** Galen M Goldscheitter, Mulugeta Seneshaw, Faridoddin Mirshahi, Evan G Buettmann, Damian C Genetos, Arun J Sanyal, Henry J Donahue

## Abstract

Metabolic Dysfunction-Associated Steatotic Liver Disease (MASLD) is highly prevalent with major risk of progression to Metabolic Dysfunction-Associated Steatohepatitis (MASH) and Hepatocellular Carcinoma (HCC). Recently, osteoporosis and bone fracture have emerged as sexually-dimorphic comorbidities of MASLD yet the mechanisms of this bone loss are unknown. Herein, we address these knowledge gaps using DIAMOND mice which develop MASLD, MASH, and HCC via Western diet exposure. We examined the skeletal phenotype of male DIAMOND mice after 16, 36, and 48 weeks of exposure to Western or control diet. At 16 weeks, male DIAMOND mice with MASLD lose trabecular bone but retain mechanical bone integrity. At 48 weeks, males lose cortical bone and mechanical integrity, indicating severe skeletal weakening. Female DIAMOND mice were protected from cortical and trabecular MASLD-associated bone loss and skeletal fragility at all timepoints. Using NicheNet, a publicly available database of hepatic mRNA expression in DIAMOND mice, and a PTH-induced model of bone loss, we suggest *Ctgf, Rarres2, Anxa2, Fgf21,* and *Mmp13* are liver-secreted ligands inducing bone resorption. This study is the first preclinical investigation of bone loss in MASLD, and the first to suggest the role of *Ctgf, Rarrest2, Anxa2, Fgf21,* and *Mmp13* as drivers of this pathology.

## INTRODUCTION

Metabolic Dysfunction-Associated Steatotic Liver Disease (MASLD) affects ∼30% of the global population and associates with increased osteoporosis and fractures. MASLD has no cure and its incidence is rapidly increasing. In ∼25% of persons with MASLD, the disease progresses to MASH (metabolic dysfunction-associated steatohepatitis), cirrhosis, chronic liver failure, and hepatocellular carcinoma, each of which associate with osteoporosis, fracture, and post-fracture mortality.[1]

Osteoporosis and fracture risk are necessary considerations in MASLD management, as people with MASLD are more likely to develop osteoporosis and more likely to experience fracture.[2], [3], [4] Further, Mendelian randomization studies identify a causal link between genetically-predicted MASLD, osteoporosis, and fracture.[5], [6] MASLD-associated fractures likely drive increased morbidity, mortality, and substantial healthcare expenditure. Effective management strategies for skeletal fragility in MASLD are urgently needed; however, the mechanisms driving this pathology remain poorly understood. As a result, therapeutic approaches are likely to remain suboptimal and lack precision until the underlying causes are elucidated. Identifying potential management strategies for bone loss in MASLD is critical to avert such undesirable outcomes.

The liver contributes to skeletal integrity through myriad well-defined mechanisms. These include, but are not limited to, its role in energy and biomolecule metabolism, vitamin D_3_ 25-hydroxylation, insulin-like growth factor-1 (IGF-1) synthesis, and sex hormone binding globulin (SHBG) synthesis.[7] In MASLD, these processes become disrupted, leading to impaired skeletal health. Moreover, MASLD associates with elevated levels of circulating inflammatory cytokine levels, including TNF, IL-6, IL-17, IFNγ, RANKL, all of which associate with or cause bone loss.[8], [9], [10] However, the exact roles of these processes in MASLD-related bone loss are yet undescribed, with one notable exception: Denosumab, an OPG-Fc mimetic that inhibits RANKL binding to RANK, improves both hepatic and skeletal phenotypes in MASLD.[11] This suggests bidirectional liver-bone crosstalk via the RANK-RANKL-OPG axis. However, denosumab indications are limited to pre-existing osteoporosis and cancerous bone lesions, it is impractically expensive as a prevention strategy, its effectiveness declines with long-term use, and rebound bone resorption after discontinuation may render it ineffective in this setting.[12] Novel prevention strategies are needed, and a suitable preclinical model is required to advance mechanistic understanding of liver-bone interactions in MASLD to this end.

Pre-clinical elaboration of skeletal consequences of MASLD/MASH is hampered by the lack of an animal model that mirrors the disease presentation and polygenic risk factors in humans. The DIAMOND (Diet-Induced Animal Model of Non-alcoholic fatty liver Disease) mouse develops liver disease solely due to high-carbohydrate, high-fat “Western” diet consumption, avoiding confounding variables of other models including micronutrient/amino acid modified diets and single gene polymorphisms.[13] DIAMOND mice rank highly among preclinical MASLD models in their transcriptomic and histologic signature relative to the human disease state.[14] The DIAMOND mouse is an isogenic cross between C57BL/6J and 129S1/SvImJ strains. Like humans, DIAMOND mice develop hallmarks of MASLD phenotype such as obesity, insulin resistance, hypertriglyceridemia, and hypercholesterolemia on a Western diet. These mice also develop hepatic steatosis after Western diet exposure of 4-8 weeks (MASLD-like) and steatohepatitis at 16-24 weeks (MASH-like). Cirrhosis and hepatocellular carcinoma (HCC) arise spontaneously by 48 weeks (**Fig. 1**).[13] DIAMOND mice are useful for the study of tissues other than liver: Nucera et al. have demonstrated their utility in the study of extrahepatic consequences of MASLD.[15] Thus, we seek to leverage DIAMOND mice to close the liver-bone knowledge gap in MASLD.

**Figure 1:**
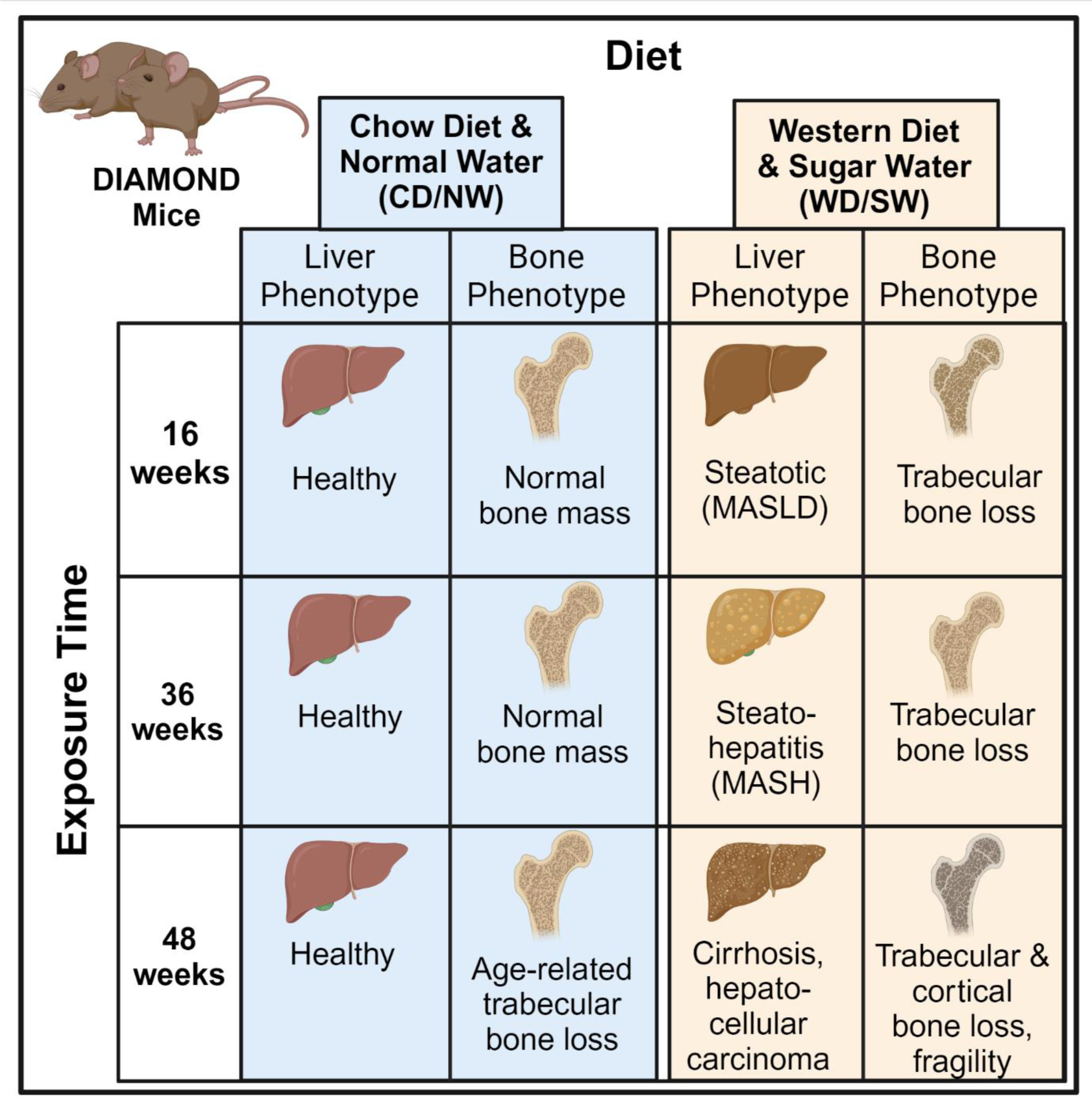
Reported skeletal and hepatic phenotypes of DIAMOND mice on chow diet & normal water versus Western diet & sugar water at 16, 36, and 48 weeks.

Clinically, MASLD affects females and males at similar rates, but bone and liver phenotypes differ by sex.[16], [17] Sex hormone levels at least partly explain these differences. Estrogens inhibit the resorption of bone by osteoclasts.[18] In the liver, estrogen influences hepatic fat deposition with evident sexual dimorphism and is implicated as protective against MASLD progression.[17] Physiologic—and elevated—levels of estrogen, therefore, are considered protective against both bone loss and MASLD. In this study, we describe the skeletal phenotype of female and male DIAMOND mice. DIAMOND mice recapitulate the timeline, circumstances, and clinical features of MASLD development in humans. Thus, we hypothesize DIAMOND mice develop sexually-dimorphic skeletal fragility alongside MASLD driven by liver-induced upregulation of pro-resorptive pathways in bone. In this work we describe the skeletal phenotype of DIAMOND mice with MASLD and identify probable molecular pathways from publicly available bone and liver RNAseq data representing treatable targets to offset the impact of this pathology. Our results suggest that changes in *Ctgf (Ccn2), Fgf21, Anxa2,* and *Mmp13* expression are involved in skeletal dysfunction associated with MASLD, which should be evaluated mechanistically in future studies.

## METHODS

### Experimental Animals

All animal care and use were overseen and approved by the Virginia Commonwealth University IACUC. Cadaveric specimens from DIAMOND mice, a well-established preclinical model of MASLD, (an isogenic cross between C57BL/6J and 129S1/SvImJ mice) were obtained from previous studies but this is the first report of skeletal effects in the DIAMOND model.[13] For our study, DIAMOND mice were randomized to a high-fat Western Diet (Teklad 88137, 42% calories from fat) and high-fructose-glucose Sugar Water (23.1 g/dL d-fructose, 18.9 g/dL d-glucose) (WD/SW) or a standard Chow Diet (Teklad 7012, 17% calories from fat) with Normal Water (CD/NW) at 8 weeks of age. Mice were humanely euthanized after 16, 36, or 48 weeks of diet exposure according to VCU protocol. Due to design of the original source studies, female DIAMOND mice were only available at 48 weeks. Hindlimbs from the carcasses were stored at -80°C after tissue isolation. As the tissues were not fixed or snap-frozen in the original design, our analysis is limited to morphologic and mechanical properties of hindlimb skeleton. The tibiae were isolated and used for micro-computed tomography and mechanical testing.

### Micro Computed Tomography

Tibiae from DIAMOND mice were embedded in 1% agarose and imaged on a Bruker SkyScan 1276 desktop micro-computed tomography scanner. The scanning parameters were 730 ms exposure time, 60 kVp voltage, 200 µA generator current, 0.5 mm aluminum filter, with an isotropic voxel resolution of 10 µm. Datasets were reconstructed in NRecon (Bruker) with parameters set to 20% beam hardening, σ=2 smoothing, and 100 ring artifact reduction. The bones were aligned along the mechanical testing support sites using DataViewer (Bruker). Cortical bone in the diaphysis and trabecular bone in the epiphysis were analyzed in CtAn (Bruker) as previously described.[19], [20] Briefly, a 180 µm segment of cortical bone was selected at the midpoint between the proximal epiphyseal plate and the distal tibiofibular junction, corresponding to the point of greatest curvature. This region was automatically segmented and thresholded at a value of 140 (8-bit pixel intensity), near the predicted site of breaking in 3-point bending. A 400 µm long epiphyseal bone region was selected immediately proximal to the epiphyseal plate. Trabecular bone in the images was manually contoured, and thresholded to a value of 120 (8-bit pixel intensity).

### Bone mechanical properties evaluation

Tibiae from DIAMOND mice were loaded to failure by breaking in 3-point bending using a Bose ElectroForce 3200 (TA Instruments, New Castle, DE). Data were captured via a 100 lbf load cell at 10 Hz with a loading rate of 1 mm/min. Tibiae were placed on supports with a span of 10 mm, loading the anteromedial surface in tension. Load was applied at the midpoint of the support span, coinciding with the point of maximal curvature of the bone. Load, deformation, stress, strain, toughness, and work were measured directly or inferred using micro-CT data, as previously described.[19], [21], [22]

### Ligand-target gene pair prediction

Ligand-target interactions in MASLD skeletal fragility were predicted in NicheNet using two publicly available datasets.[23] This approach has been used to evaluate crosstalk between the nervous and skeletal systems, but our work is the first to apply it to the liver and skeleton.[24] The first dataset, GSE67678, contains hepatic tissue of male DIAMOND mice fed a high-fat diet and high-fructose-glucose solution (n=5) or control diet (n=5) for 8 weeks; this dataset was defined as the ligand donor for ligand-target interactions. Because no sequencing data for skeletal tissue gene expression in DIAMOND mice exists, a model of continuous PTH administration in osteoblasts was selected as a highly catabolic state of skeletal metabolism. The target dataset, from Li *et al.* (S*upp Table 4* <u>and *5*</u>), identified genes in cortical bone which were regulated by continuous PTH(1-34) (cPTH) in rats.[25] 3-month old female Sprague-Dawley rats received 4 μg/100 g/day PTH(1-34) via implantable osmotic pump. DEGs were defined as those with a fold change (log_2_FC) ≥ 1 and adjusted p-value < 0.05. Ligand-receptor interactions were predicted using NicheNet based on downstream effectors altered in response to continuous PTH. Predicted ligand-target interactions in MASLD skeletal fragility were generated using NicheNet.[23] Hepatic ligands were identified by filtering differentially expressed genes (DEGs) in mice fed WD/SW compared to those fed CD/NW, which were integrated to modeled ligand-gene target pairs. Similarly, skeletal receptors and affected genes included only those predicted to be regulated by the identified hepatic ligands by the NicheNet prior knowledge model. Regulatory potential, interaction potential, and ligand activity were ranked by area under the precision recall curve (AUPR).

### Statistical Analysis

Equal variances and normality were assessed using the Bartlett and Shapiro-Wilk tests, respectively. Due to normality and equal variances, inter-group differences were assessed via two-way ANOVA. Post-hoc analyses were conducted using Tukey’s method. Multiple comparisons were controlled via the Bonferroni method. Ligand-target gene assessments were conducted using NicheNet as described above.[23] Sample numbers were determined by sample availability. Among males, 5 mice were available per group at 16 weeks of diet exposure, 10 mice per group at 36 weeks, and 10 mice per group at 48 weeks. Among females, 5 mice per group were available at 48 weeks. Ideally, 8-12 mice would be employed per group for a study of this nature. Applicable exclusion criteria were the presence of a pre-existing fracture (pre- or post-mortem).

## RESULTS

### Male DIAMOND mice with MASLD develop skeletal fragility and lose bone in a time- and bone compartment-dependent manner

Compared to CD/NW controls, male DIAMOND mice on WD/SW lost bone, evident in trabeculae at 16 weeks (**Figure 2A**, **Table 1**) and cortices at 48 weeks (**Figure 2C**, **Table 1**). Trabecular bone volume fraction, trabecular number, and trabecular spacing exhibit their most severe deleterious effects at 16 weeks of WD/SW exposure compared to CD/NW (**Figure 2B**, **Table 1**). Trabecular thickness remains largely unaltered by exposure to WD/SW (**Figure 2B**, **Table 1**). The trabecular bone phenotype among male DIAMOND mice on WD/SW and CD/NW is similar at 36- and 48-weeks of diet exposure.

**Figure 2:**
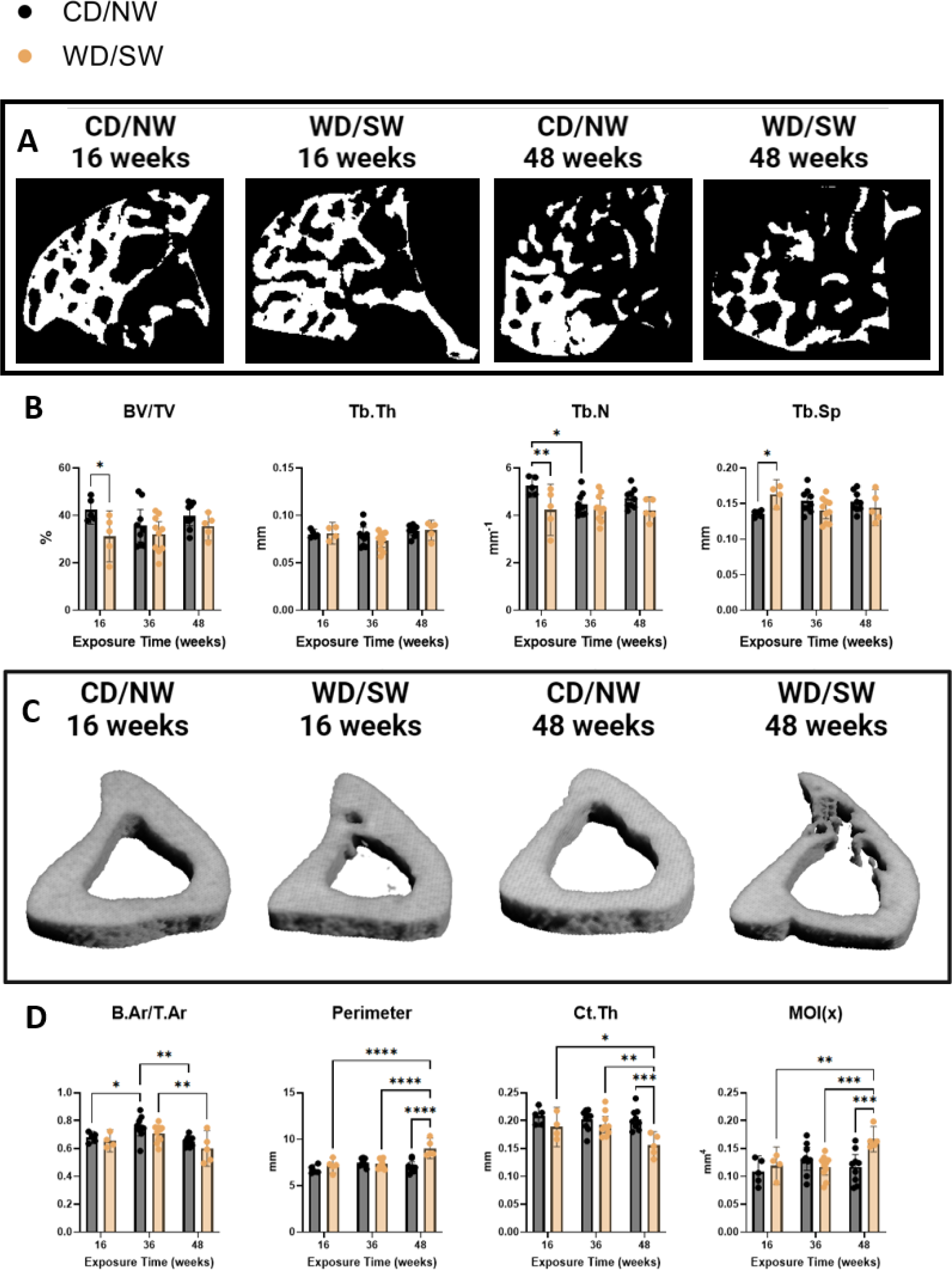
Bone morphometry of Male DIAMOND mice at 16, 36, and 48 weeks of CD/NW or WD/SW exposure. **(A)** 2D projections of epiphyseal trabecular bone. **(B)** Proximal epiphyseal trabecular bone volume/tissue volume (BV/TV), trabecular thickness (Tb.Th), trabecular number (Tb.N), and trabecular spacing (Tb.Sp). **(C)** 3D projections of the tibial mid-diaphysis. **(D)** Mid-diaphyseal cortical bone area/tissue area (B.Ar/T.Ar), perimeter, cortical thickness (Ct.Th), and moment of inertia in the x-direction (MOI(x)). (* p < 0.05, ** p < 0.01, ***, p < 0.001, **** p < 0.0001)

**Table 1:**
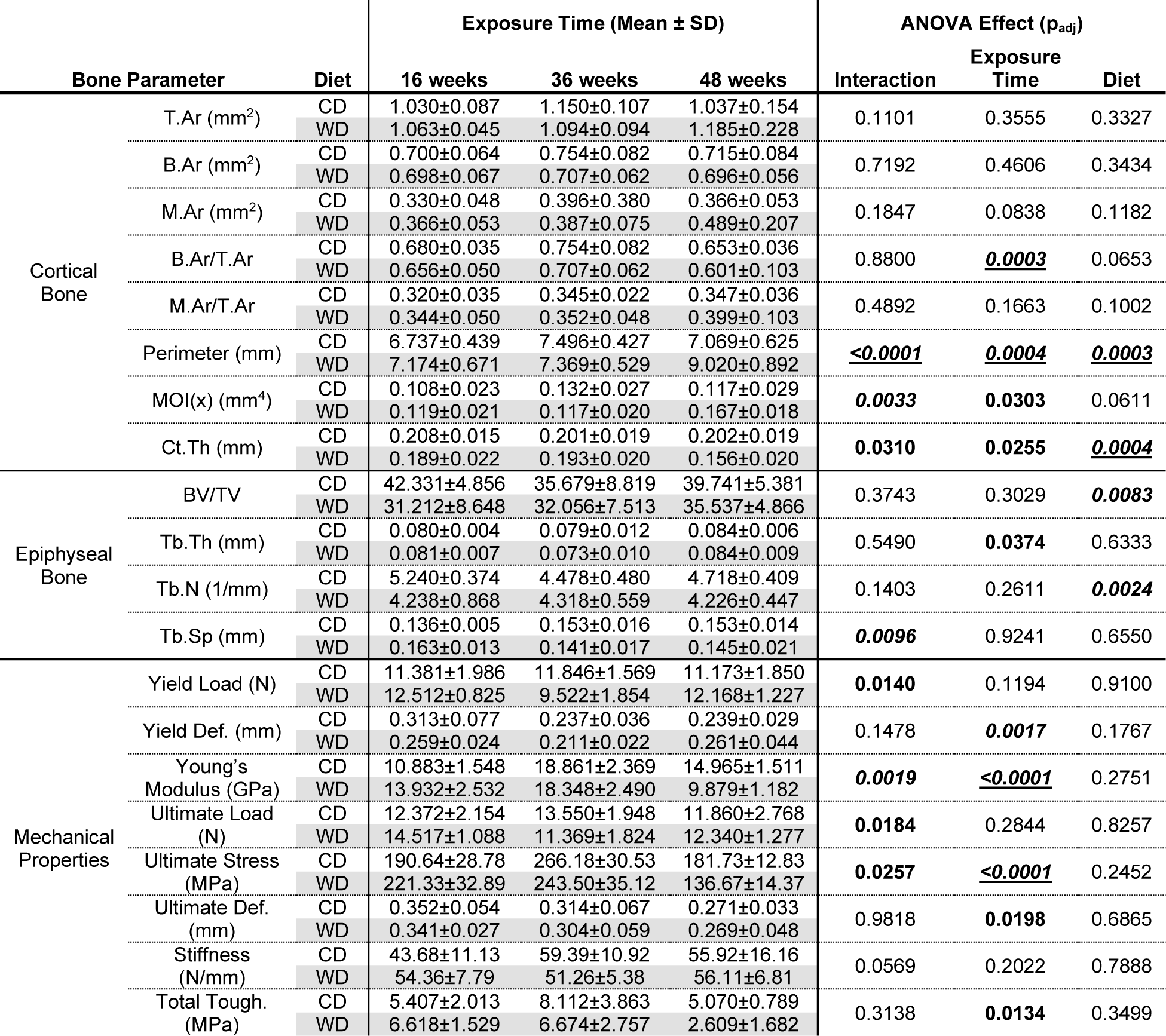
Geometric and mechanical parameters from male DIAMOND mice fed CD/NW or WD/SW for 16, 36, or 48 weeks beginning at 8 weeks of age. (**Bold:** p < 0.05, ***bold + italics:*** p < 0.01, ***<u>bold + italics + underline:</u>***p < 0.001)

Within the mid-diaphysis of the tibia, cortical thickness decreased markedly after 48 weeks of WD/SW exposure compared to CD/NW in male DIAMOND mice (**Figure 2D**, **Table 1**). This is reflected in significant increases in cortical perimeter and moment of inertia (**Figure 2D**, **Table 1**). Bone area fraction is largely conserved (**Figure 2D**, **Table 1**). In summary, after 48 weeks of diet exposure, cortical bone in WD/SW mice thinned and increased in diameter, likely reducing fracture resistance.

Yield stress, ultimate stress, Young’s modulus, and total toughness decrease among male DIAMOND mice with MASLD compared to those without in diet- and exposure time-dependent manners (**Figure 3A, 3B, Table 1**). At 16 weeks, their mechanical properties are similar, but mice with MASLD progressively develop fragility during diet exposure. In summary, WD/SW exposure in the male MASLD DIAMOND mouse model associate with early, deleterious trabecular changes that are minimally observable at late disease stages, and a pronounced decline in cortical bone parameters and mechanical integrity at late disease stages.

**Figure 3:**
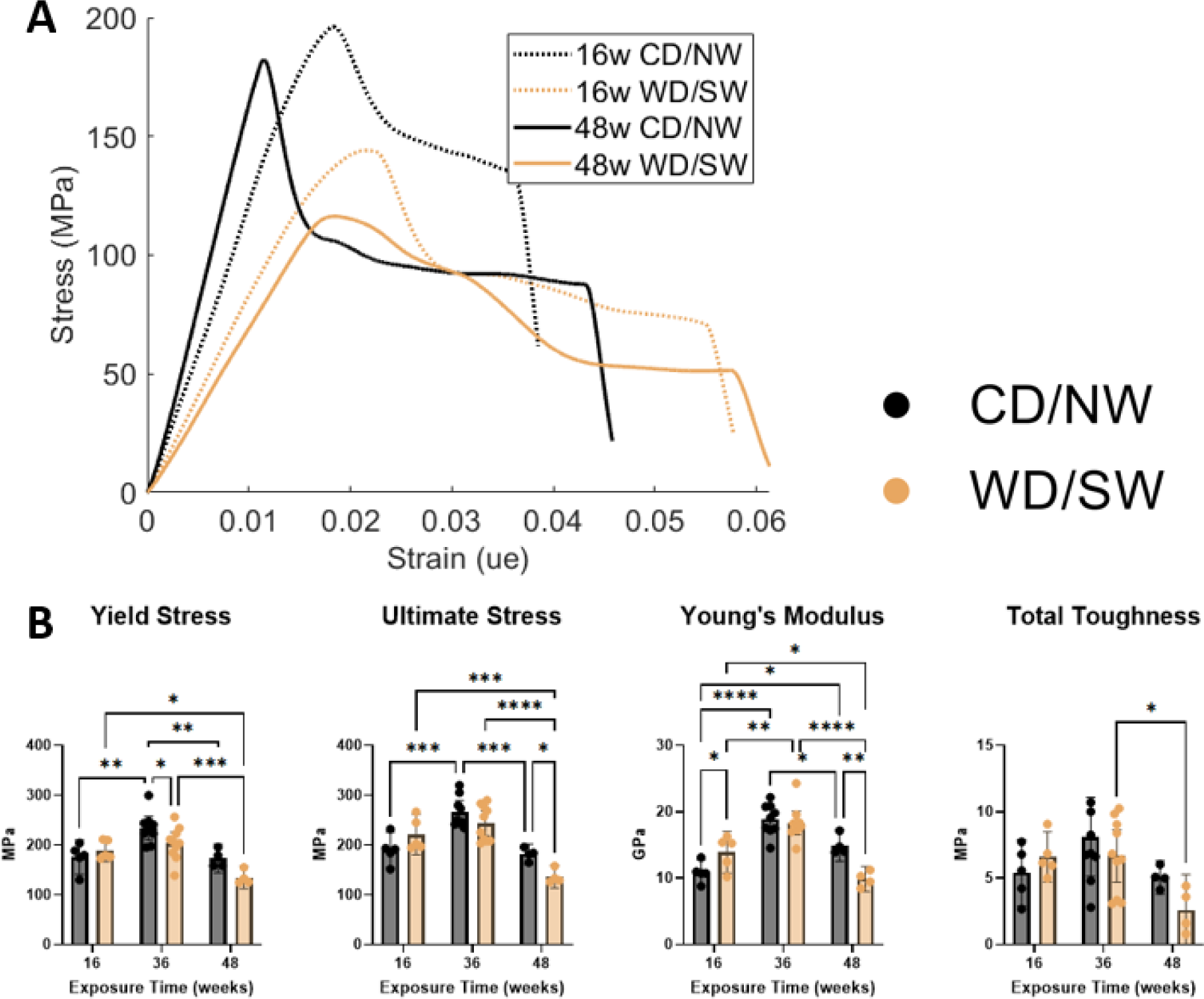
Bone mechanical properties of male DIAMOND mice at 16, 36, and 48 weeks of CD/NW or WD/SW exposure. **(A)** Representative 3-point bending traces from DIAMOND mice tibias. **(B)** Tibial yield stress, ultimate stress, young’s modulus, and total toughness derived from 3-point bending. (MOI(x)). (* p < 0.05, ** p < 0.01, ***, p < 0.001, **** p < 0.0001)

### Bone loss in DIAMOND mice with MASLD is sexually dimorphic

Inclusion of female DIAMOND mice with MASLD at 48 weeks, showed sex-based differences in the bone phenotype, with females partially protected in epiphyseal and diaphyseal bone properties compared to males. While males with MASLD demonstrated losses in bone area fraction, cortical thickness, and ultimate stress before failure, compared to same-sex CD/NW controls, females maintained their skeletal geometry and mechanical integrity in all these indices. Females had greater epiphyseal bone volume fraction and trabecular number, and had lower trabecular spacing, than males while on WD/SW. (**Figure 4A, 4B, Table 2**). Female DIAMOND mice gained bone area fraction and cortical thickness while on WD/SW, whereas males experienced losses. (**Figure 4C, 4D, Table 2**) Further, male DIAMOND mice with MASLD experienced increases in cortical perimeter and moment of inertia while females did not (**Figure 4D**, **Table 2**).

**Figure 4:**
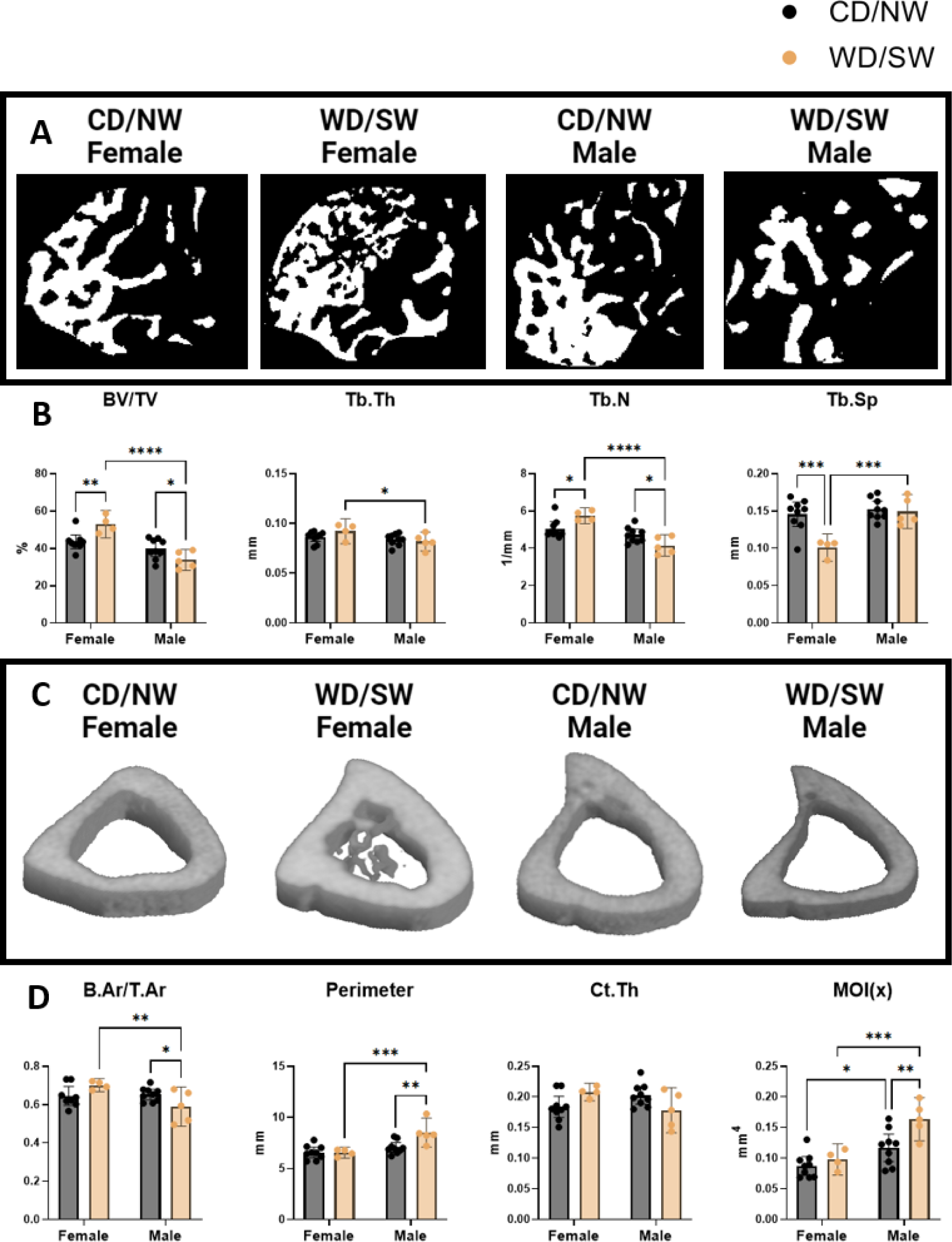
Sexual dimorphism of bone morphology among female and male DIAMOND mice after 48 weeks of CD/NW or WD/SW exposure from micro-CT. **(A)** 2D projections of epiphyseal trabecular bone. **(B)** Proximal epiphyseal trabecular bone volume/tissue volume (BV/TV), trabecular thickness (Tb.Th), trabecular number (Tb.N), and trabecular spacing (Tb.Sp). **(C)** 3D projections of the tibial mid-diaphysis. **(D)** Mid-diaphyseal cortical bone area/tissue area (B.Ar/T.Ar), perimeter, cortical thickness (Ct.Th), and moment of inertia in the x-direction (MOI(x)). (* p < 0.05, ** p < 0.01, ***, p < 0.001, **** p < 0.0001)

**Table 2:**
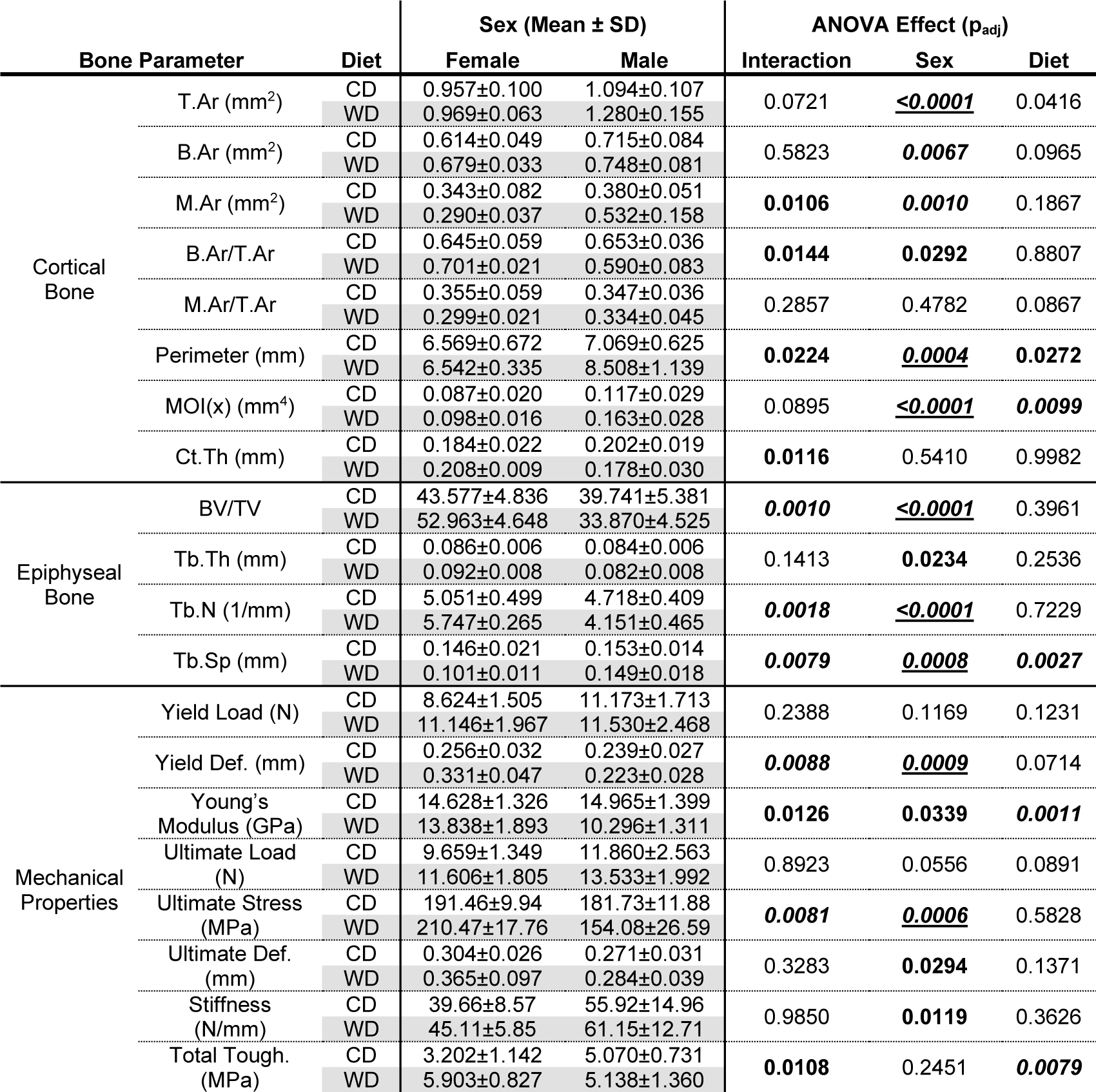
Geometric and mechanical parameters from male and female DIAMOND mice fed CD/NW or WD/SW for 48 weeks beginning at 8 weeks of age. (**Bold:** p < 0.05, ***bold + italics:*** p < 0.01, ***<u>bold + italics + underline:</u>*** p < 0.001)

The sexually-dimorphic impact of MASLD on skeletal microarchitecture extended to mechanical properties of tibiae. Female DIAMOND mice given WD/SW do not exhibit losses in yield stress, ultimate stress, or Young’s Modulus compared to same-sex controls as seen in male DIAMOND mice, which exhibit an increase in total toughness after 48 weeks of WD/SW exposure vs CD/NW (**Figure 5B**, **Table 2**). Marked cortical thinning was observed in males on WD/SW, but not in males on CD/NW or females on either diet (**Figure 4A**, **Table 2**). In summary, these data suggest female DIAMOND mice are protected from the deleterious skeletal changes observed among males when consuming WD/SW vs CD/NW for at least 48 weeks.

**Figure 5:**
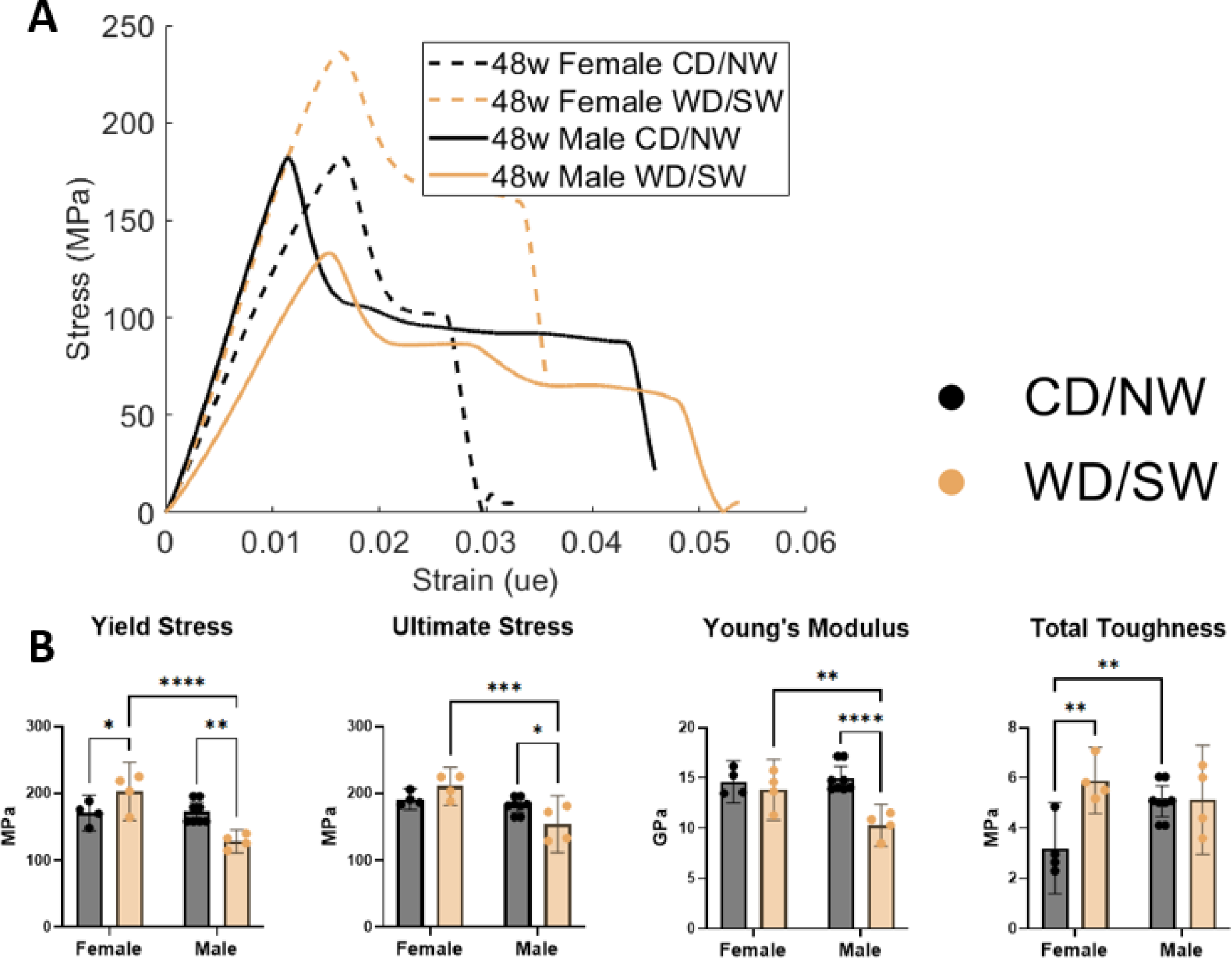
Sexual dimorphism of bone mechanical properties among female and male DIAMOND mice after 48 weeks of CD/NW or WD/SW exposure. **(A)** Representative 3-point bending traces from DIAMOND mice tibias. **(B)** Tibial yield stress, ultimate stress, young’s modulus, and total toughness derived from 3-point bending. (MOI(x)). (* p < 0.05, ** p < 0.01, ***, p < 0.001, **** p < 0.0001)

### *In silico* analysis of liver-bone crosstalk reveals probable molecular pathways driving bone loss in MASLD

Considering the shared pathways by which MASLD and bone loss arise, it is highly probable skeletal fragility in MASLD is driven by liver-bone crosstalk. Our approach used NicheNet to describe hepatoskeletal crosstalk in MASLD **(Figure 6A**) NicheNet differential regulation analysis on publicly available hepatic RNASeq data from DIAMOND mice (**Figure 6B**)[13] versus bone cell receptor activity prediction based on cPTH-induced changes in bone[25] identified multiple hepatic ligands (encoded by genes *Ccn2, Rarres2, Anxa2, Apoc1, Apoe,* among others) with prioritized impact on bone gene expression *(Ccnd1, Postn, Aebp1,* among others). DIAMOND mice fed WD/SW vs CD/NW show a dramatically different liver transcriptomic expression profile, clustering neatly via UMAP dimension reduction (**Figure 6A**). Among these hepatic genes, we select meaningfully and significantly upregulated genes with a secreted isoform (**Figure 6B**). The top 25 of these, ranked by ligand potential, are shown in **Figure 6D**, where *Ccn2, Rarres, Anxa2, Apoc1, and Apoe* having the highest probability of possessing ligand activity on receptors in bone. **Figure 6E** shows interaction potential, derived from the NicheNet prior learning model[23], between secreted hepatic ligands in **Figure 6D** and expressed cognate receptors in bone, showing biological plausibility these ligands affect downstream effector pathways in bone. Finally, the regulatory potential of the identified set of hepatic ligands on the top 18 downstream effector genes in bone is shown in **Figure 6C**. Among these, hepatocyte-secreted *Ccn2* is predicted to induce upregulation of *Ccnd1* in bone is the interaction with the greatest regulatory potential. Notably, *Fgf21* from hepatocytes is expected to induce upregulation of numerous genes in bone, including *Ccnd1, Postn, Alpl,* and *Lox*. The comprehensive integration of ligand activity, regulatory potential, and receptor expression profiles enhances our understanding of the molecular mechanisms driving gene expression changes in response to a skeletally-catabolic stimulus to hepatic dysfunction.

**Figure 6:**
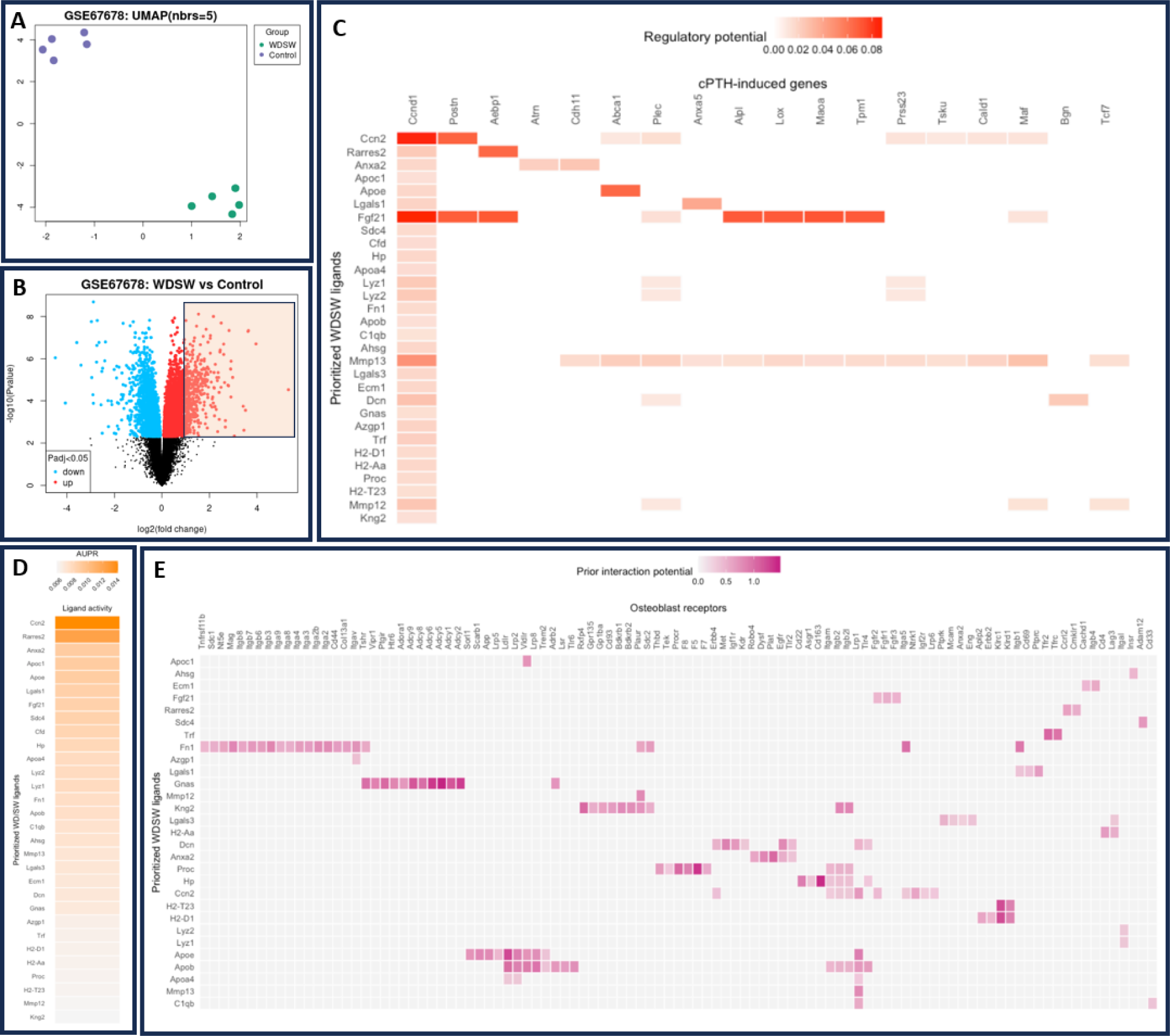
NicheNet-predicted ligand-affected gene interactions between 8-week old DIAMOND mice with MASLD and osteoblasts from Sprague-Dawley rats receiving a highly-catabolic cPTH dosing regime. **(A)** clustering of hepatic gene expression by UMAP. **(B)** volcano plot showing hepatic differentially expressed genes (DEGs) among DIAMOND mice with MASLD vs healthy controls. Colored box indicates DEGs with log_2_(FC) > 1 and -log_10_(p_adj_) > 2, indicating the DEGs from which ligands are selected. **(C)** regulatory potential of ligands (selected from the DEGs shown in **B**) from WD/SW mice, and the bone genes they are predicted to induce in a state of rapid bone catabolism. **(D)** Potential ligand activity of DEGs from DIAMOND mice with MASLD compared to healthy controls. **(E)** Hepatic ligands (from **B**) and their cognate receptors in bone cells, with heatmap intensity indicating the interaction potential of each ligand with its cognate receptor.

## DISCUSSION

The DIAMOND mouse is a well-established preclinical model of MASLD. DIAMOND mice develop MASLD, MASH, cirrhosis, and HCC on a high-fat, high-carbohydrate diet alone, a result of a polygenic inheritance pattern. These features in addition to changes in histologic disease severity, serum metabolic profile, and inflammatory serum milieu–make the DIAMOND mouse an appropriate preclinical candidate for the study of extrahepatic effects of MASLD, as demonstrated in hypothalamic metabolism. In this work, we are the first to observe and report, sex- and duration of diet-dependent changes in skeletal fragility among DIAMOND mice with MASLD.

Among male DIAMOND mice, MASLD drives myriad changes in the appendicular skeleton. In its early stages, MASLD is associated with reductions in trabecular bone parameters. However, as diet exposure time lengthens, trabecular bone differences between mice with MASLD are much smaller compared to those without MASLD. Beyond 16 weeks of diet exposure, mice with MASLD lose minimal trabecular bone beyond what is already lost, suggesting rapid trabecular bone loss between weeks 0 and 16 which then stabilizes. Meanwhile, mice without MASLD eventually mirror the trabecular bone phenotype of mice with MASLD as—what are presumably age-related—changes accumulate. By 48 weeks, trabecular bone of mice with and without MASLD are nearly identical. Indeed, trabecular bone volume fraction in the tibia of C57BL6 mice is expected to peak at or before 2 months of age, and monotonically decrease throughout their remaining lifespan.[26] The same pattern of growth and resorption is presumably present in DIAMOND mice; however, the expected date of peak trabecular bone mass is unknown due to its mixed C57BL/6J and 129S1/SvIm background. By 16 weeks, DIAMOND mice will have developed MASLD and may experience early MASH. The same inflammatory state driving hepatic disease development likely causes trabecular bone resorption in the skeleton. Trabecular bone in male DIAMOND mice on WD/SW experiences deleterious changes in early diet exposure yet remains stable from weeks 16 to 48.

In the cortical bone compartment, differences between male mice fed CD/NW and WD/SW are not apparent until 48 weeks of diet exposure suggesting slower accumulation of deleterious changes. This is reflected in losses in mechanical integrity measured in 3-point bending, in which most variance is explained by cortical bone morphology and tissue properties, especially cortical thickness. Additionally, unlike most structural indices, mechanical properties do not exhibit monotonic decline during the study period. The ultimate stress and Young’s modulus rise from 16 to 36 weeks, then fall from 36 to 48 weeks. This is consistent with age-related changes to cortical bone, as mice commonly reach peak cortical thickness and mechanical integrity at 6 months of age or later. In our study, the 16-week timepoint occurs at 24 weeks of age, as mice are randomized to their assigned diet at 8 weeks of age. C57BL/6 mice—a founder strain of the DIAMOND mouse—typically reach peak cortical thickness at 6 months of age.[26] There are no data regarding 129S1/SvIm mice—the other DIAMOND founder strain—or DIAMOND mice describing the age at which they achieve peak skeletal integrity. Our data suggest peak skeletal integrity occurs in DIAMOND mice sometime between 16 and 48 weeks of diet exposure (or 24 and 56 weeks of age), likely around 36 weeks of diet exposure (44 weeks of age). Interestingly, the only timepoint with a substantial difference in ultimate load between the CD/NW and WD/SW groups occurred at 36 weeks. The greatest change in ultimate load was observed at 36 weeks, while cortical bone area fraction, cortical thickness, and ultimate stress are most affected at 48 weeks. It should be noted that hepatocellular carcinoma remains a confounder, as DIAMOND mice will frequently develop spontaneous hepatocellular carcinoma by 48 weeks of age and should be considered in future analyses.

Female DIAMOND mice appear protected against MASLD-associated skeletal fragility at 48 weeks of WD/SW exposure vs CD/NW. The presence of increased estrogens in female compared to male mice[27] is likely to be protective against MASLD-associated bone loss because estrogen is, independently, protective against the progression of both MASLD and bone loss.[16], [17] Whether this phenotype is observed in trabecular bone could not be assessed as female DIAMOND mice at 16 weeks of diet exposure were not available for analysis. Further, RNASeq data for female DIAMOND mice is also unavailable. As such, our analysis of potential hepatic ligand and affected skeletal genes could only be conducted in males. The mechanisms by which female DIAMOND mice are protected against bone cannot be addressed in this work, but are the subject of ongoing studies. Female DIAMOND mice are less affected by both MASLD and its associated bone loss when exposed to identical conditions as males. Their histologic disease, degree of hepatomegaly, tumor burden, and skeletal fragility are less severe than those of male DIAMOND mice. There is, therefore, a correlative association between sex, MASLD severity, and skeletal fragility. Although not directly measured here, sex differences in eating habits, hyperglycemia, insulin sensitivity, and hormonal signaling of DIAMOND mice remain unknown, as well as differences in the serum proteome, hepatic and skeletal transcriptome. Each of these components likely plays a role in the development of both MASLD and skeletal fragility and may be responsible for its sexual dimorphism. These components must be addressed in future studies to elaborate the mechanism of protection from MASLD and bone loss among female DIAMOND mice.

As this study was conducted *a posteriori*, important confounding metrics of metabolism and skeletal health were not measured as covariates. Prospective studies in this area would benefit by measuring total caloric intake and expenditure during the study period and total activity level. While DIAMOND mice do not exhibit hyperglycemia or insulin resistance at 16 or 36 weeks of diet exposure, they exhibit both at 48 weeks.[13] As such, they should be assessed in future studies of bone loss in MASLD.

Male DIAMOND mice exposed to WD/SW after 48 weeks demonstrated a catabolic skeletal phenotype, as evidenced by decreased bone microarchitecture and strength. To infer potential mechanisms for increased skeletal fragility, NicheNet analysis was performed on publicly available RNAseq liver data in male DIAMOND mice[13] and a separate bone gene set[25] undergoing a catabolic state induced by cPTH treatment. Among the 25 highest regulatory potential ligands identified in our analysis, *Ctgf* (*Ccn2*)*, Rarres2*, *Anxa2*, *Fgf21*, and *Mmp13* have biological plausibility to modulate bone metabolism under cPTH treatment and may be mechanistic drivers of MASLD driven skeletal fragility. For example, *Ctgf* (*Ccn2*) has variable effects on bone the impact of which is dictated by developmental stage and interactions with other competing or cooperating local signals. *Ctgf* is a regulator of normal skeletal morphology during development.[28] However, when overexpressed in the adult skeleton, *Ctgf* induces bone loss.[29] The role of hepatic-derived *Ctgf* on skeleton function is heretofore unconsidered; our data suggest increased hepatic *Ctgf* expression in livers characterized by MASLD, drives excessive bone turnover and net loss. *Rarres2*, which encodes the adipokine chemerin, is associated with bone loss in both humans and mice.[30], [31], [32] Chemerin is highly expressed in hepatocytes, and its serum levels are increased in persons with MASLD[33] and MASH.[34] Chemerin has been shown to induce osteoclastogenesis and inhibit osteoblastogenesis *in vitro*.[30] Chemerin is, therefore, a probable link between MASLD, MASH, and bone loss in humans and mice. *Anxa2* expression, which encodes Annexin A2, is associated with fragility fracture and the development of osteoporosis in humans.[35], [36] Decreased osteoblast formation and increased membrane-bound RANKL synthesis are proposed as mechanisms for this effect.[36], [37] Our data suggest hepatic *Anxa2* is a potential mediator of MASLD-associated bone loss, via its induction of *Atrn* (encodes attractin) and *Cdh11* (encodes cadherin 11) in osteoblasts. *Fgf21* is a regulator of glucose and lipid metabolism, driving increased insulin sensitivity and decreased serum glucose and triglycerides.[38], [39] Systemic administration of Fgf-21 also corrects obesity in diet-induced and ob/ob mice.[40] As such, *Fgf21* has been proposed as a promising drug for metabolic diseases, which would include MASLD. However, *Fgf21* overexpression drives substantial bone loss in mice and its withdrawal promotes a high bone-mass phenotype.[41] Thus, *Fgf21* overexpression—and purported ligand activity—, make it a probable candidate driving bone loss in male DIAMOND mice with MASLD. Lastly, *Mmp13* encodes for a matrix metalloprotease, highly expressed in osteoblasts and critical for collagen reorganization during bone mineralization. It is a drug development target in osteoarthritis therapy, where it has been identified as a mediator of bone destruction around the articular surfaces.[42] Further, its expression in breast cancer bone metastases drives osteolysis and osteoclastogenesis.[43] Deletion of *Mmp13* in mesenchymal cells increases bone mass and may attenuate bone loss associated with estrogen withdrawal.[44] While the role of hepatic expression of *Mmp13* in skeletal fragility has not been elaborated, we propose its ligand activity increases bone loss by inducing osteoclast-mediated bone resorption, which could be explored in future conditional genetic studies targeting *Mmp13* in the liver of DIAMOND mice.

Bone RNASeq data is not yet available from DIAMOND mice. As such, we selected a highly-catabolic cPTH regime as our target dataset for NicheNet affected gene predictions.[25] This approach has several shortcomings. First, the skeletal mechanisms of bone loss in the target dataset may be distinct from those in MASLD. Second, the identity of the target cells contained within the population is not characterized, and is likely a mixture of osteocytes, osteoblasts, osteoclasts, bone lining cells, bone marrow stromal cells, vascular endothelial cells, and hematogenous cell populations. Third, cPTH is substantially more rapidly catabolic than the effects of MASLD-associated bone loss. Therefore, identification of the involvement of *Ctgf* (*Ccn2*), *Rarres2*, *Anxa2, Fgf21*, and *Mmp13* in MASLD associated bone loss requires further validation in bone RNAseq data from MASLD mice. Nonetheless, the engagement of *Ctgf* (*Ccn2*), *Rarres2, Anxa2, Fgf21*, and *Mmp13* in our current study, which are heavily implicated in bone metabolism, validates our approach using NicheNet in further specific datasets to our bone phenotype in MASLD.

Skeletal fragility in MASLD is an emerging complication of a highly prevalent, incurable metabolic disorder. Given MASLD affects roughly one quarter of the global population, the implications of increased rates of fracture, hospitalization, and early mortality are immense. Additionally, these are likely to drastically accelerate healthcare spending given an aging, increasingly overweight/obese population. Identifying mechanisms driving this pathology is critical. We propose *Ctgf (Ccn2), Rarres2, Anxa2, Fgf21, and Mmp13* encode novel, plausible hepatic ligands driving bone loss in MASLD, based on computational NicheNet analysis from liver RNAseq data, as targets for future mechanistic study.

## CONCLUSION

Bone loss in MASLD is an important consideration in the management of this disease. In this study, we observed trabecular and cortical bone loss—at different time points—in DIAMOND mice. Thus, we propose DIAMOND mice to be an excellent candidate for the study of this combined hepato-skeletal pathology. The DIAMOND mouse mimics the hepatic phenotype of humans with MASLD and has already been used in the study of extrahepatic manifestations of the disease. We observe congruency between the sexual dimorphism in skeletal phenotype, marked by skeletal deterioration primarily in male DIAMOND mice on the diet for 48 weeks herein, and identified elsewhere in humans with MASLD. Further, we identify putative ligands of hepato-skeletal crosstalk, including *Ctgf* (*Ccn2*), *Fgf21*, *Anxa2*, and *Mmp13*. These ligands are associated with low bone mass in mice, osteoporosis, and fragility fracture. and therefore strong putative mediators of bone loss in MASLD, which should be studied in future preclinical models of MASLD before evaluation as therapeutic targets.

